# *Drosophila* Males Use 5′-to-3′ Phased Biogenesis to Make *Stellate*-silencing piRNAs that Lack Homology to Maternally Deposited piRNA Guides

**DOI:** 10.1101/2022.09.12.507655

**Authors:** Zsolt G. Venkei, Ildar Gainetdinov, Margaret R. Starostik, Charlotte P. Choi, Peiwei Chen, Chiraag Balsara, Troy W. Whitfield, George W. Bell, Suhua Feng, Steven E. Jacobsen, Alexei A. Aravin, John K. Kim, Philip D. Zamore, Yukiko M. Yamashita

**Affiliations:** Whitehead Institute for Biomedical Research, Department of Biology, Massachusetts Institute of Technology, Cambridge, MA, U.S.A.; RNA Therapeutics Institute, University of Massachusetts Medical School, Worcester, MA, U.S.A.; Department of Biology, Johns Hopkins University, Baltimore, MD, U.S.A.; Division of Biology and Biological Engineering, California Institute of Technology, Pasadena, CA, U.S.A.; University of Michigan, Life Sciences Institute, MI, U.S.A.; Department of Molecular, Cell and Developmental Biology, University of California, Los Angeles, PO Box 957239, Los Angeles, CA 90095-7239, USA; Eli and Edyth Broad Center of Regenerative Medicine and Stem Cell Research, University of California, Los Angeles, Los Angeles, CA 90095, USA; Howard Hughes Medical Institute, Los Angeles, CA 90095, USA; Howard Hughes Medical Institute, University of Massachusetts Chan Medical School, Worcester, MA, U.S.A.; Howard Hughes Medical Institute, Whitehead Institute, Cambridge, MA, U.S.A.

## Abstract

PIWI-interacting RNAs (piRNAs) direct PIWI proteins to silence complementary targets such as transposons. In animals with a maternally specified germline, e.g. *Drosophila melanogaster*, maternally deposited piRNAs initiate piRNA biogenesis in the progeny. Normal fertility in *D. melanogaster* males requires repression of tandemly repeated *Stellate* genes by piRNAs from *Suppressor of Stellate* [*Su(Ste)*]. Because the *Su(Ste)* loci are on the Y chromosome, *Su(Ste)* piRNAs are not deposited in oocytes. How the male germline produces *Su(Ste)* piRNAs in the absence of maternally deposited *Su(Ste)* piRNAs is unknown. Here, we show that *Su(Ste)* piRNAs are made in the early male germline via 5′-to-3′ phased piRNA biogenesis triggered by maternally deposited *1360/Hoppel* transposon piRNAs. Strikingly, deposition of *Su(Ste)* piRNAs from XXY mothers obviates the need for phased piRNA biogenesis in sons. Together, our study uncovers the developmentally programmed mechanism that allows fly mothers to protect their sons using a Y-linked piRNA locus.

## Introduction

The PIWI-interacting RNA (piRNA) pathway is an animal-specific, small RNA-mediated mechanism that silences transposable elements and other selfish genetic elements (Ozata et al. 2019). Loss of piRNAs reduces fertility due to derepression of TEs (Girard et al. 2006; Aravin et al. 2007; Brennecke et al. 2007; Das et al. 2008), or deregulation of gene expression (Wu et al. 2020; Chen et al. 2021a; Choi et al. 2021). At the core of piRNA-mediated target silencing are 18-35-nt piRNAs that bind to and guide PIWI proteins to their targets via nucleotide sequence complementarity (Aravin et al. 2006; Girard et al. 2006; Grivna et al. 2006; Lau et al. 2006; Vagin et al. 2006). The three *Drosophila melanogaster* PIWI proteins have specialized functions in the germline: Piwi represses transposon transcription in the nucleus, whereas cytoplasmic Ago3 and Aubergine (Aub) cleave piRNA precursor and transposon transcripts in the cytoplasm (Pal-Bhadra et al. 2004; Saito et al. 2006; Vagin et al. 2006; Brennecke et al. 2007; Sienski et al. 2012; Huang et al. 2014; Le Thomas et al. 2014; Post et al. 2014; Han et al. 2015; Mohn et al. 2015; Senti et al. 2015; Wang et al. 2015).

To make new piRNAs, animals use pre-existing piRNAs to direct slicing of complementary transcripts, initiating piRNA biogenesis from cleavage products (Gainetdinov et al. 2018). For example, in the *D. melanogaster* female germline, Ago3 and Aub are loaded with piRNAs derived from complementary transcripts (transposon mRNAs and piRNA precursors), and the 3′ cleavage product of Ago3 slicing is used to make antisense Aub-loaded piRNAs and vice versa. This positive feedback loop known as the ‘ping-pong’ cycle amplifies the transposon-targeting population of piRNAs (Brennecke et al. 2007; Gunawardane et al. 2007). The ping-pong pathway also initiates 5′-to-3′ fragmentation of the remainder of the cleavage product into tail-to-head, phased piRNAs loaded in Piwi (Han et al. 2015; Homolka et al. 2015; Mohn et al. 2015; Yang et al. 2016). This process is carried out by the endonuclease Zucchini (Zuc; PLD6 in mammals) with the help of the RNA helicase Armitage (Armi; MOV10L1 in mammals) (Pane et al. 2007; Ge et al. 2019; Munafo et al. 2019; Yamashiro et al. 2020).

The ping-pong cycle requires pre-existing piRNAs to initiate the amplification process. In *D. melanogaster*, maternally deposited piRNAs serve this purpose, providing the first pool of piRNAs that can initiate the ping-pong cycle (Blumenstiel and Hartl 2005; Brennecke et al. 2008; de Vanssay et al. 2012; Le Thomas et al. 2014; de Albuquerque et al. 2015). For example, the inability of mothers to provide P-element-derived piRNAs in a cross between naïve mothers and P-element-infested fathers causes derepression of selfish elements, leading to sterile offspring, a phenomenon called hybrid dysgenesis (Kidwell et al. 1973; Kidwell and Kidwell 1976; Kidwell et al. 1977; Ronsseray et al. 1984; Brennecke et al. 2008; Khurana et al. 2011; Teixeira et al. 2017; Wakisaka et al. 2017; Moon et al. 2018; Srivastav et al. 2019).

*Stellate (Ste)* and *Suppressor of Stellate* [*Su(Ste)*] in *D. melanogaster* provided the founding paradigm of piRNA-directed repression in *D. melanogaster* (Hardy et al. 1984; Livak 1984; McKee and Satter 1996; Kalmykova et al. 1998; Belloni et al. 2002). *Ste* is a repetitive gene whose unchecked expression results in the formation of Ste protein crystals, an amyloid-like protein aggregate that causes male sterility via unknown mechanisms (Bozzetti et al. 1995). To ensure male fertility, *Ste* genes on the X chromosome are normally repressed by *Su(Ste)* piRNAs that are antisense to *Ste* and are produced from the Y chromosome (Aravin et al. 2001; Aravin et al. 2003; Aravin et al. 2004; Vagin et al. 2006). *Su(Ste)* locus is composed of tandem repeats that have high level (~90%) of identity to *Ste* sequence. *Ste* is the major silencing target of the piRNA pathway in the *D. melanogaster* male germline (Aravin et al. 2001; Aravin et al. 2003; Nishida et al. 2007; Nagao et al. 2010; Quenerch′du et al. 2016; Chen et al. 2021a), requiring *armi, zuc, aub* and *ago3*, but not *piwi* or *rhino* (*rhi*), suggesting that *Ste* repression is primarily dependent on cytoplasmic cleavage of the *Ste* mRNA (Vagin et al. 2006; Pane et al. 2007; Klattenhoff et al. 2009; Chen et al. 2021b). Because *Su(Ste)* is encoded on the Y chromosome, fly mothers— which lack a Y chromosome—cannot provide their sons with *Su(Ste)* piRNAs to initiate biogenesis. How then is *Ste* repressed in the apparent absence of maternally deposited piRNAs?

Here, we describe the mechanism by which the male germline represses *Ste* in the absence of maternally deposited *Su(Ste)* piRNAs. We show that *Su(Ste)* piRNAs are produced by an Armi- and Zuc-dependent phased piRNA biogenesis in male germline stem cells (GSCs) and early spermatogonia (SGs), days before expression of the *Ste* target in the spermatocytes. Phased biogenesis of *Su(Ste)* piRNAs in GSCs/SGs is critical to repress *Ste* later in spermatocytes, and thus for male fertility. XX mothers cannot deposit Y-linked *Su(Ste)* piRNAs to their sons. Instead, our data show that males from XX mothers utilize maternally deposited *1360/Hoppel* piRNAs to cleave *Su(Ste)* precursors and initiate 5′-to-3′ phased biogenesis of *Su(Ste)* piRNAs in the early germline (GSCs/SGs). We show that the requirement for Armi, a protein essential for phased piRNA biogenesis, in *Su(Ste)* piRNA production in males is relieved when XXY females provide maternal *Su(Ste)* piRNAs to their sons’ germline. These data explain how maternally deposited piRNAs can direct production of non-homologous piRNA guides in the germline of the progeny. Our study reveals a mechanism for intergenerational transmission of piRNA-coded memory in the absence of direct homology that can protect offspring from selfish genetic elements not encountered by their mothers.

## Results

### Transcription of *Su(Ste)* piRNA Precursors Starts in Germline Stem Cells, Days Before *Ste* Expression

To investigate *Su(Ste)* piRNA precursor expression and processing into piRNAs during *D. melanogaster* spermatogenesis, we used single-molecule RNA fluorescent *in situ* hybridization (smRNA-FISH) (Raj and Tyagi 2010; Fingerhut et al. 2019). By leveraging single nucleotide polymorphisms between *Ste* and *Su(Ste)*, we used a single *in situ* probe and a collection of Stellaris *in situ* probes to specifically visualize *Su(Ste)* and *Ste*, respectively (see Methods). We detected expression of *Su(Ste)* piRNA precursor transcripts only from the genomic strand that produces transcripts antisense to *Ste* mRNAs (not shown). smRNA-FISH can detect *Ste* mRNAs and *Su(Ste)* precursor transcripts but not mature piRNAs, because small RNAs are not retained in formaldehyde-fixed tissues.

smFISH revealed that in wild-type testes, *Ste* transcripts are first detected in the nuclei of spermatocytes (Figure 1A, B, F). In contrast, in the absence of *Su(Ste)* in XO males, *Ste* transcripts were readily detected in the spermatocyte cytoplasm (Figure 1C, G), leading to production of Ste protein crystals, a known cause of subfertility. Notably, in XO males, cytoplasmic *Ste* mRNA was only observed in spermatocytes (Figure 1C), suggesting that *Ste* is transcriptionally silent in early germ cells (i.e. GSCs and SGs) (Figure 1C, E). Our smRNA-FISH experiments readily detected *Su(Ste)* expression in GSCs, earlier than previously reported (Aravin et al. 2004). Thus, *Su(Ste)* expression precedes that of *Ste* by ~2-3 days. The steady-state abundance of nuclear *Su(Ste)* transcripts peaked in late SGs/early spermatocytes and was undetectable by the time *Ste* expression was first detected, in late spermatocytes (Figure 1B, D, F).

**Figure 1.**
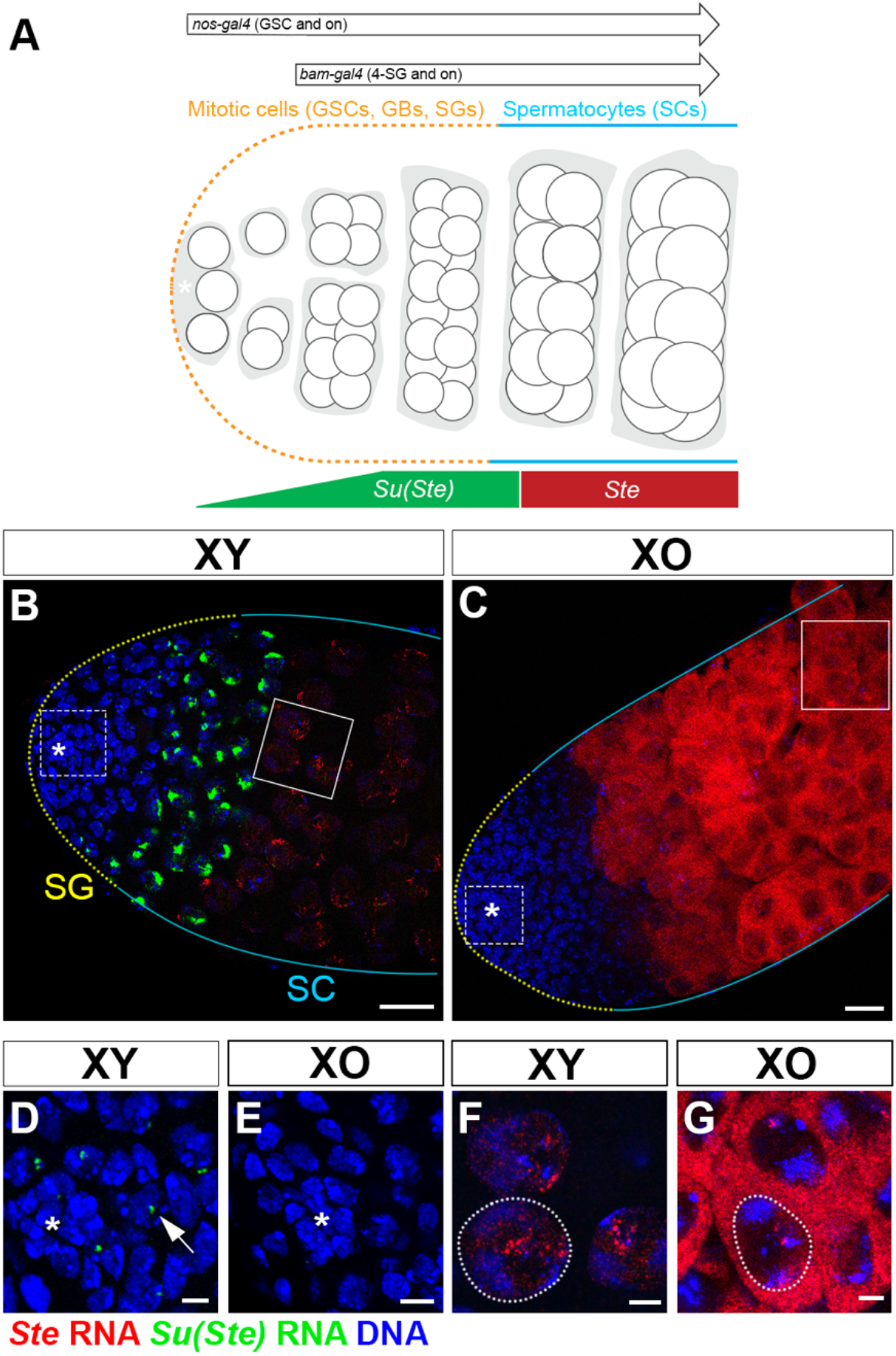
*Su(Ste)* transcription precedes that of *Ste* during germ cell differentiation. (A) Early stages of *D. melanogaster* spermatogenesis. The stem cell niche is formed by non-dividing somatic cells (hub, marked by asterisk). The germline stem cells (GSCs) are physically attached to the hub, and divide asymmetrically. The gonialblasts (GBs), the differentiating daughters of GSCs, undergo four rounds of mitotic divisions with incomplete cytokinesis. Resultant 16-cell spermatogonia (SGs) then enter meiotic prophase as spermatocytes. The expression patterns of *nos-gal4* and *bam-gal4* drivers in the adult male germ line are also indicated. GSCs/early SGs are indicated by yellow dotted line, zone of spermatocytes by cyan lines in this and all subsequent figures. (B-C) Expression of *Ste* (red) and *Su(Ste)* (green) in the wild type (B) and in XO (C) testes. (D-E) Inserts from B and C magnifying the stem cell niche in wild type (D) and XO (E) testes. Arrow points to *Su(Ste)* transcripts in a GSC nucleus. (F-G) Magnified inserts from B and C show subcellular *Ste* localization in wild type (F) and XO (G) spermatocytes. Dotted white lines indicate the nuclear periphery. Hub (*), *Ste* RNA (red), *Su(Ste)* RNA (green), DAPI (blue). Bars 20 μm for B-C, and 5 μm for D-G.

Ping-pong amplification of *Ste*-targeting piRNAs should require the presence of both *Su(Ste)* and *Ste* RNA in the same cells. Our data suggest that ping-pong amplification is unlikely to explain the biogenesis of *Su(Ste)* piRNAs, because *Su(Ste)* piRNA precursors are transcribed and processed into *Ste*-targeting piRNAs before the first detectable accumulation of *Ste* mRNA.

### *zuc*- and *armi*-Dependent Processing of *Su(Ste)* piRNA Precursor Transcripts in Germline Stem Cells and Spermatogonia

We find that processing of *Su(Ste)* precursors into mature piRNAs in GSCs/SGs depends on components of the phased piRNA biogenesis pathway. In wild type GSCs/SGs, *Su(Ste)* transcripts were detected as a single nuclear focus, corresponding to nascent transcripts from the *Su(Ste)* loci (Figure 2A, F). In contrast, in *armi^1/72.1^* or *zuc^EY11457/-^* loss-of-function mutants, the nuclear foci of *Su(Ste)* transcripts were enlarged and multiple cytoplasmic foci appeared, likely representing accumulation of unprocessed piRNA precursor transcripts (Figure 2C, E, H, K). Similar *Su(Ste)* cytoplasmic foci were detected when *armi* or *zuc* mRNA was specifically depleted in germ cells by RNAi using pVALIUM22 transgenes *(armi^TRIP.GL00254^* and *zuc^TRIP.GL00111^*; henceforth, *armi^RNAi^* and *zuc^RNAi^)* driven by *nanos(nos)-Gal4* (Van Doren et al. 1998) (Figure 1A). The appearance of *Su(Ste)* cytoplasmic foci in *zuc* and *armi* mutants (Figure 2K) is consistent with the increase in the steady-state abundance of *Su(Ste)* transcripts measured by RT-qPCR in *zuc^EY11457/-^* mutant testis enriched for SGs by over-expressing *dpp* (Figure S1).

**Figure 2.**
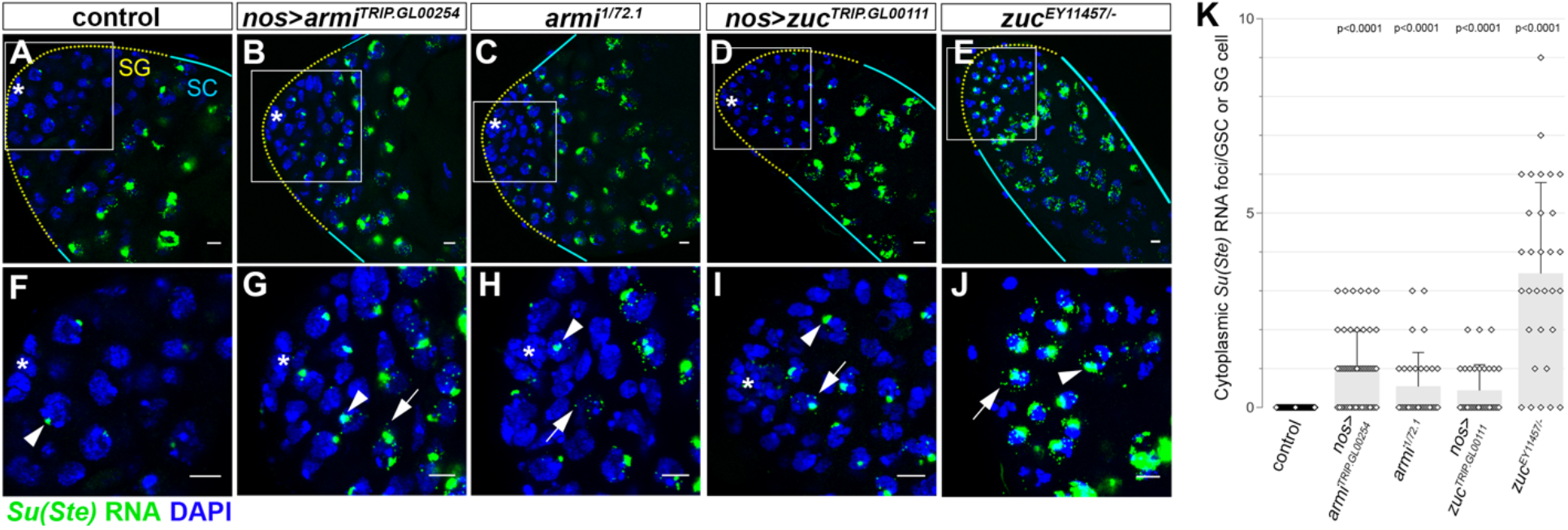
*Su(Ste)* precursor transcripts are increased in GSCs and SGs of *armi* and *zuc* mutant testes. *Su(Ste)* transcript (green) in wild type testis (A, F), and in piRNA pathway mutant testes of the indicated genotypes (B-E, G-J). (F-J) magnified regions of the niche from A-E (marked by quadrates). Loss-of-function allelic combinations *armi^1/72.1^* and *zuc^EY1145/-7^* were used. RNAi constructs *(armi^TRIP.GL00254^, zuc^TRIP.GL00111^)* were expressed by *nos-gal4*, which drives expression in early germ cells (GSCs and onward). GSC/early SGs are indicated by yellow dotted line, zone of spermatocytes by cyan lines. Arrowheads point to nuclear transcripts, arrows point to cytoplasmic RNA foci. Hub (*), DAPI (blue), bars 5 μm. K) Quantification of cytoplasmic *Su(Ste)* RNA foci in GCSs and SG cells. Data are presented as mean±s.d. *N*=30-90 nuclei/genotype. P-value from Welch′s unequal variances t-test (unpaired, two-tailed) is provided compared to control.

By contrast, *Su(Ste)* piRNA precursor transcripts did not accumulate when *aub, ago3*, or *vas* mRNAs were depleted by *nos*-driven RNAi (Figure S2). Because Aub, Ago3, and Vasa are required for ping-pong amplification of piRNAs, these results suggest that in GSC/SGs the production of piRNAs from *Su(Ste)* transcripts is dominated by the phased piRNA biogenesis pathway.

### *Ste* Silencing Requires *zuc* and *armi* in Early Male Germ Cells

Repression of *Ste* in late spermatocytes depends on *zuc* and *armi* expression during a short window in early spermatogenesis. When *armi* or *zuc* mRNA was depleted by *nos*-driven RNAi *(nos>armi^RNAi^* or *nos>zuc^RNAi^)* throughout the germline (Figure 1A), we observed derepression of *Ste* RNA (Figure 3A-C, G), accompanied by Ste protein accumulation (Figure 3I) and reduced fertility (Figure 3J). In contrast, using *bam-gal4* (Figure 1A) to deplete *armi* or *zuc* in >4-cell SG stages *(bam>armi^RNAi^* or *bam>zuc^RNAi^)* had no observable effect on *Ste* repression or fertility (Figure 3D, H, I, J), suggesting that *armi* and *zuc* are dispensable for *Ste* repression after the four-cell SG stage.

**Figure 3.**
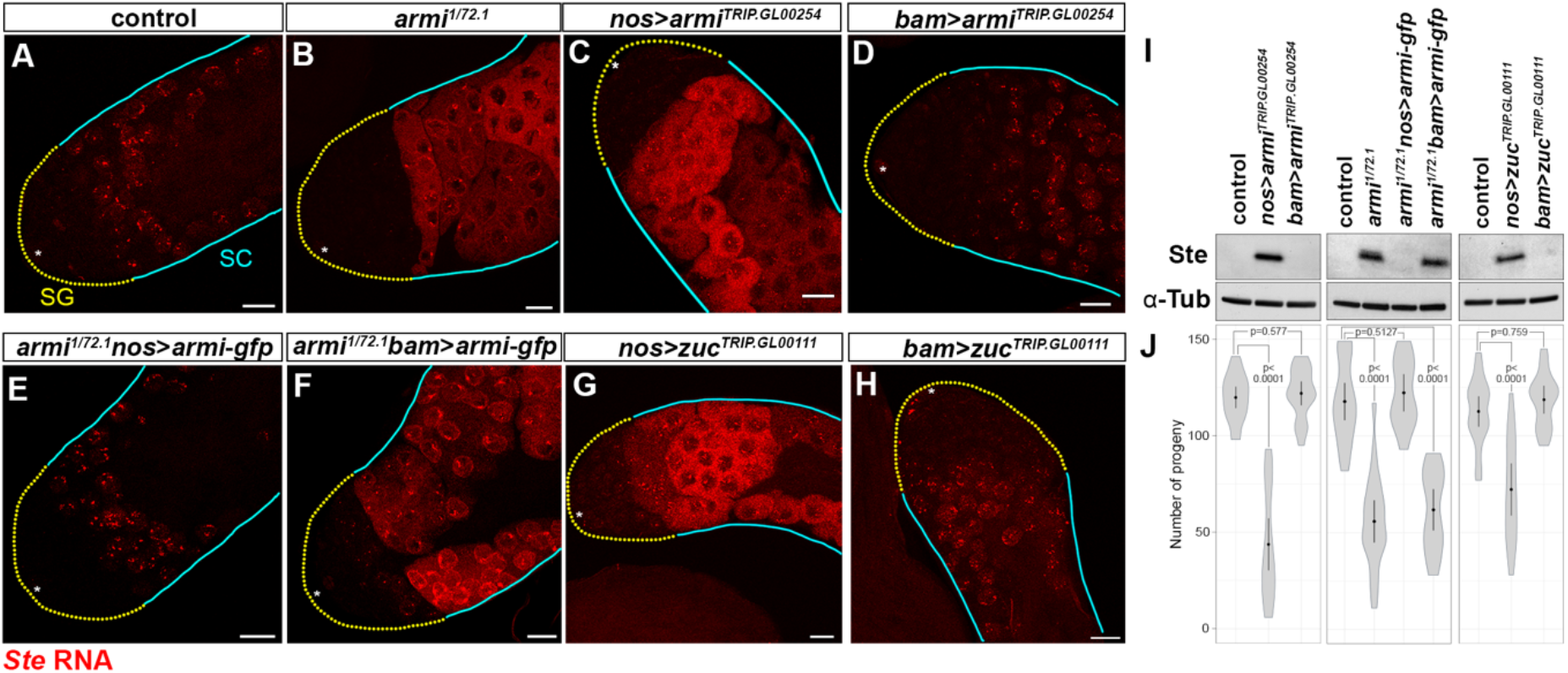
*armi* and *zuc* are specifically required in GSCs/early SGs to repress *Ste*. *Ste* smRNA-FISH (red) in the testes from control (A), *armi^1/72.1^*, (B), *nos>armi^RNAi^* (*armi^TRIP.GL00254^*) (C), *bam>armi^RNAi^* (D), *armi^1/72.1^* expressing *armi-gfp* with *nos-gal4* (E), *armi^1/72.1^* expressing *armi-gfp* with *bam-gal4* (F), *nos>zuc^RNAi^ (zuc^TRIP-GL00111^)* (G) or *bam>zuc^RNAi^* (H). GSCs/early SGs are indicated by yellow dotted line, zone of spermatocytes by cyan lines. Hub (*), bars 20 μm. (I) Anti-Stellate and anti-Tubulin western blots of whole testis lysates from the indicated genotypes. (J) Male fertility of indicated genotypes (number of progeny/male/7 days). Data are presented as mean±s.d. *N*=20 male per genotype. P-value from Welch′s unequal variances t-test (unpaired, two-tailed) is provided compared to control.

Consistent with the idea that *Ste* silencing requires Armitage in early germ cells, expression in *armi*^1/72.1^ of an *armi-gfp* transgene under the control of *nos-gal4* restored *Ste* repression (Figure 3E, I, J). In contrast, expression of the same rescue construct but driven by *bam-gal4* failed to rescue the *armi* mutant phenotype (Figure 3F, I, J). Conversely, both *nos-gal4*- and *bam-gal4*-driven RNAi of *aub, ago3*, or *vas* led to derepression of *Ste* in spermatocytes (Figure 4A-E, H, I). Yet expressing a *gfp-aub* rescue transgene using *bam-gal4* driver essentially restored *Ste* repression in the loss-of-function *aub* mutant *(aub^HN2/QC42^)* (Figure 4F, G), demonstrating that zygotic expression of ping-pong pathway genes after the 4-cell SG stage is required to repress *Ste*.

**Figure 4.**
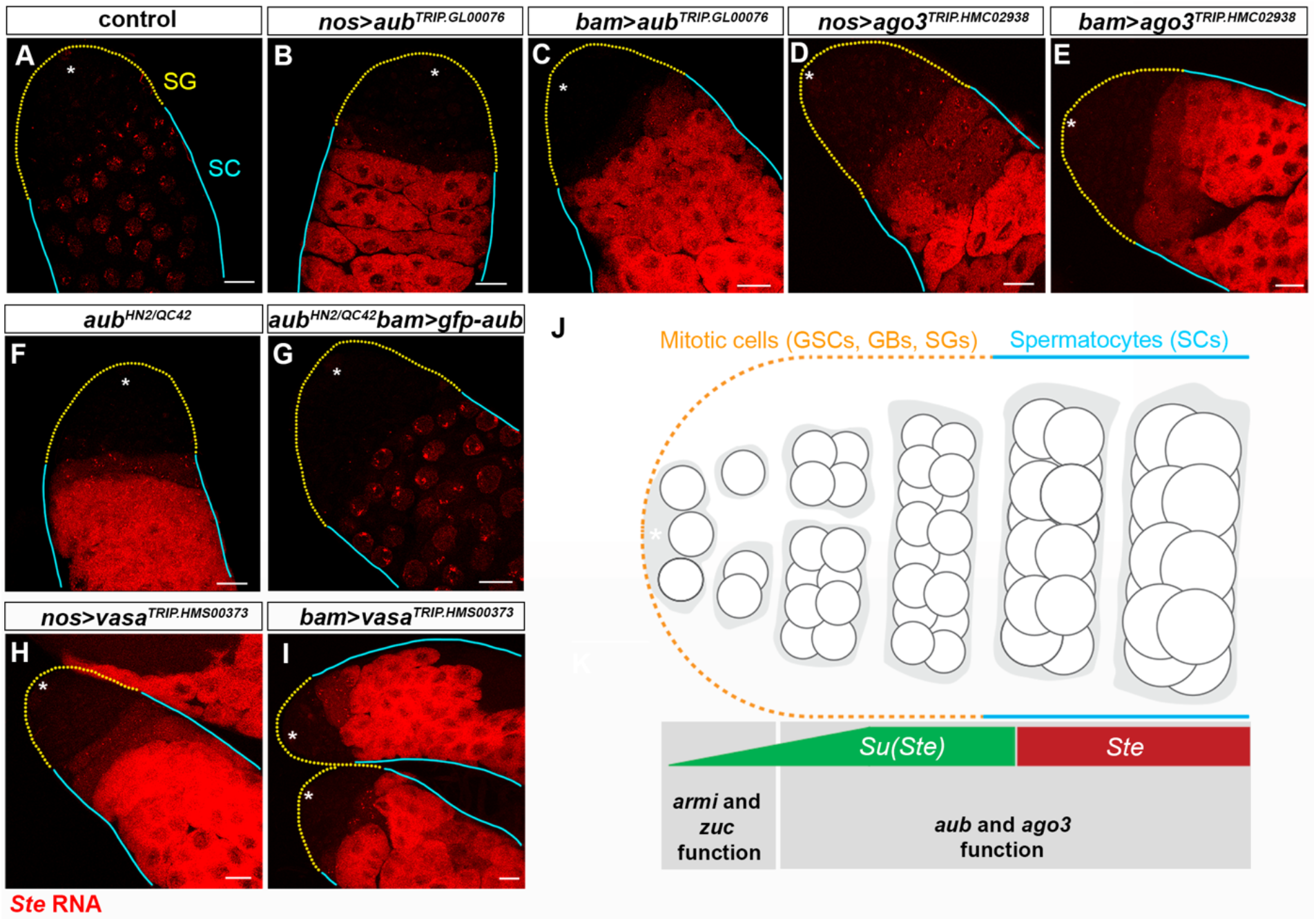
Ping-pong pathway components, *aub, ago3* and *vasa*, are specifically required in later SGs/spermatocytes to repress *Ste*. (A-I) *Ste* smRNA-FISH (red) in the testes from control (A), *nos>aub^RNAi^ (aub^TRIP.GL00076^*), (B), *bam>aub^RNAi^* (C), *nos>ago3^RNAi^ (ago3^TRIPHMC02938^)* (D), *bam> ago3^RNAi^* (E), *aub^HN2/QC42^* (F), *aub^HN2/QC42^ expressing gfp-aub* with *bam-gal4* (G), *nos>vas^RNAi^ (vas^TRIP.HMS00373^)* (H), *bam>vas^RNAi^* (I) testes. GSCs/early SGs are indicated by yellow dotted line, spermatocytes by cyan lines. Hub (*), bars 20 μm. (J) Summary of the developmental stages where *Su(Ste)* and *Ste* are expressed, and where *armi*/*zuc*, and *aub*/*ago3* are required.

We conclude that *Su(Ste)* piRNA biogenesis and piRNA-directed silencing of *Ste* are temporally separated during fly spermatogenesis: *Su(Ste)* piRNAs are produced in a *zuc*- and *armi*-dependent manner in early germ cells (GSC to four-cell SG) and repress *Ste* later in spermatocytes via Aub- and Ago3-catalyzed cleavage.

### *1360/Hoppel* piRNAs Trigger Phased Biogenesis of *Su(Ste)* piRNAs

Efficient repression of *Ste* requires production of *Su(Ste)* piRNAs days before *Ste* is first expressed (Figure 1, 4J). Production of *Su(Ste)* piRNAs in early male germ cells requires Zuc and Armi, components of the phased piRNA biogenesis pathway (Figure 2, 3, 4J). Typically, phased piRNA biogenesis is initiated by a piRNA-directed slicing event that generates a long 5′ monophosphorylated cleavage product (pre-pre-piRNA). The pre-pre-piRNA is then fragmented by Zuc into phased, tail-to-head piRNAs (Wang et al. 2014; Han et al. 2015; Homolka et al. 2015; Mohn et al. 2015). But *Ste* piRNAs that could trigger phased fragmentation of *Su(Ste)* precursors are not produced by mothers (see below).

We propose that phased production of *Su(Ste)* piRNAs is initiated by maternally inherited *1360/Hoppel* transposon-derived piRNAs that direct cleavage of the *1360/Hoppel* sequence residing at the 5′ end of *Su(Ste)* precursors (Figure 5A). Several observations support this idea: (1) transcription of *Su(Ste)* starts inside a *1360/Hoppel* transposon insertion upstream of the sequence complementary to *Ste* (Aravin et al. 2001); (2) ovaries contain abundant *1360/Hoppel* transposon-derived piRNAs (~18,200 ± 400 per 10 pg of total RNA); and (3) mothers deliver *1360/Hoppel* piRNA to their male offspring via the oocyte (Brennecke et al. 2008).

**Figure 5.**
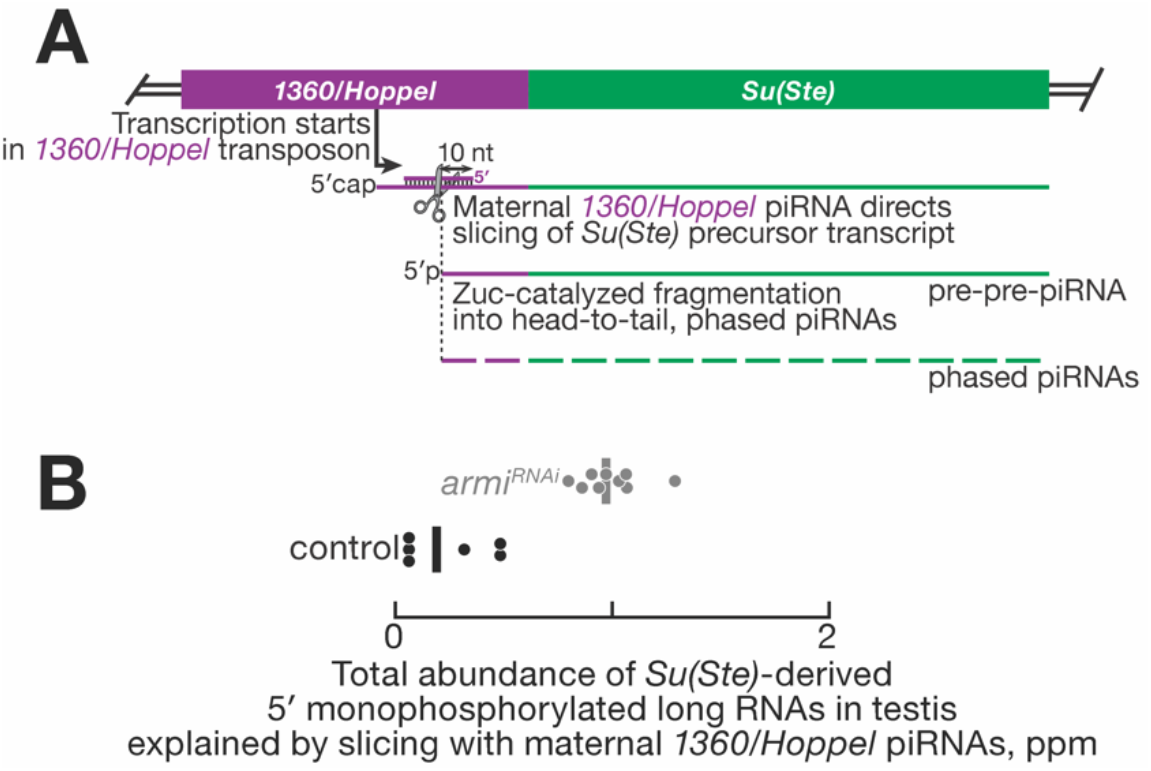
Trigger piRNAs for phased *Su(Ste)* piRNA biogenesis in males. (A) Model for initiation of phased biogenesis of *Su(Ste)* piRNAs by maternal *1360/Hoppel* piRNAs. (B) Putative pre-pre-piRNAs produced via cleavage guided by maternal 1360/Hoppel piRNAs in male gonads. Data are from all possible permutations of ovarian small RNA (*n* = 3) and 5′ monophosphorylated long RNA data sets (*n* = 2 for control; *n* = 3 for *armi^RNAi^).*

To test this model, we sequenced ≥ 200-nt long, 5′ monophosphorylated RNAs from adult testis to identify putative *Su(Ste)* pre-pre-piRNAs whose 5′ ends lie in the upstream *1360/Hoppel* insertion and are explained by piRNA-directed cleavage. Like all Argonautes, PIWI proteins cleave their targets between nucleotides t10 and t11, the target nucleotides complementary to piRNA nucleotides g10 and g11. For *Su(Ste)*-derived long RNAs overlapping both the upstream transposon insertion and the sequence complementary to *Ste*, the 5′ ends of ~40% of these long RNAs lay between nucleotides g10 and g11 of an antisense maternal *1360/Hoppel* piRNA. Supporting the idea that these long RNAs are pre-pre-piRNAs processed by phased biogenesis pathway, their steady-state abundance was ~five-fold higher when phased biogenesis in males was blocked using *nos*-driven *armi^RNAi^* (Figure 5B). Together, these data suggest that *Su(Ste)* precursor transcripts in early male germ cells are sliced by maternally deposited transposon-derived piRNAs.

### *Su(Ste)* piRNAs Made in XXY Females Silence *Ste* in the Germline of Progeny

Our model assumes that piRNA•PIWI complexes deposited by mothers can cleave complementary RNAs in the germline of their sons. To experimentally test this assumption, we used XXY female flies to artificially produce *Su(Ste)* piRNAs in oocytes. Y chromosome-encoded *Su(Ste)* piRNA precursors and *Su(Ste)* piRNAs were detected in XXY (2,700 ± 80 piRNAs per 10 pg total RNA) but not XX ovaries (30 ± 30 piRNAs per 10 pg total RNA; Figure 6A and S3). These maternally produced *Su(Ste)* piRNAs were able to repress a*gfp-Ste* transgene in XXY females (Figure S4). Strikingly, when *Su(Ste)* piRNA biogenesis was blocked in sons, maternal *Su(Ste)* piRNAs from XXY oocytes sufficed to silence *Stellate* in the testis: unlike *nos>armi^RNAi^* males from XX mothers (Figure 3I), *nos>armi^RNAi^* sons derived from XXY females effectively repressed *Ste* (Figure 6B-I and S5). We conclude that maternal deposition of *Su(Ste)* piRNAs by XXY mothers suffices to silence *Ste* mRNA and bypasses the requirement for phased piRNA production pathway in early male germ cells.

**Figure 6.**
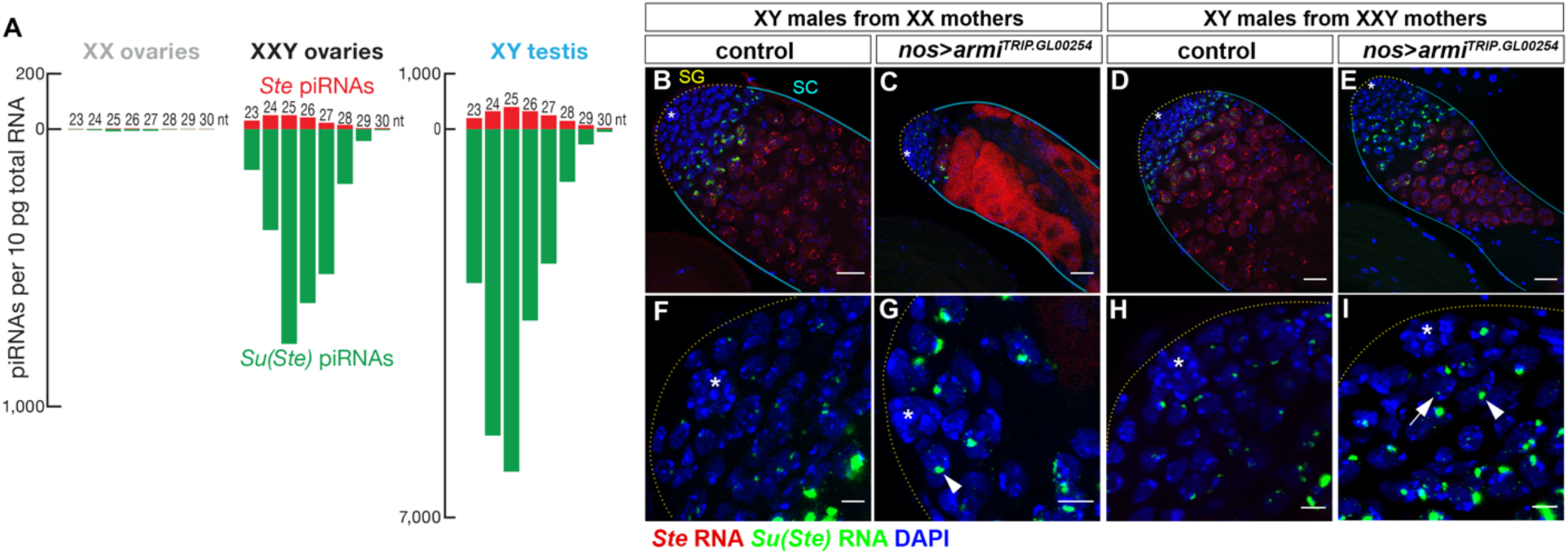
Maternally deposited *Su(Ste)* piRNAs can rescue *Ste* repression in *armi^RNAi^* male germline. (A) Length profile of *Ste*- and *Su(Ste)*-derived piRNAs in XX, XXY ovaries, and XY testis. (B-E) Testes of control (B) and *nos>armi^RNAi^* (C) sons from XX mothers, showing derepression of *Ste* in *nos>armi^RNAi^*, and testes of control (D) and *nos>armi^RNAi^* (E) sons from XXY mothers, showing that *nos>armi^RNAi^* sons can repress *Ste*, if they are from XXY mothers. *Ste* RNA (red), *Su(Ste)* piRNA precursor (green), DAPI (blue). (F-I) *Su(Ste)* piRNA precursor in situ hybridization signals at the apical tip of the testis from the indicated genotypes. Arrowheads point to enhanced nuclear transcripts, arrow points to a cytoplasmic RNA, indicating incomplete piRNA processing in *nos>armi^RNAi^.* Bars 20 μm for B-E, and 5 μm for F-I.

High-throughput sequencing of long or small RNA from GSCs/early SGs is infeasible, because early germ cells constitute a small fraction of the adult testis. Artificial expression of *Su(Ste)* precursors in XXY ovaries however allowed us to test whether the production of phased *Su(Ste)* piRNAs in XXY females is initiated by *1360/Hoppel* transposon piRNAs that direct cleavage of *Su(Ste)* transcripts. Among ≥ 200-nt long, 5′ monophosphorylated RNAs from XXY ovaries, we identified putative *Su(Ste)* pre-pre-piRNAs that could have been produced by *1360/Hoppel* piRNA-guided slicing (Figure 7A, top). When we confined our analysis to *Su(Ste)* long RNAs spanning both the *1360/Hoppel* and *Ste*-derived sequences, we found that the 5′ ends of ~35% of such long RNAs overlapped an antisense *1360/Hoppel* piRNA by exactly 10 nt, suggesting that these monophosphorylated RNAs are pre-pre-piRNAs generated by transposon piRNA-directed slicing.

**Figure 7.**
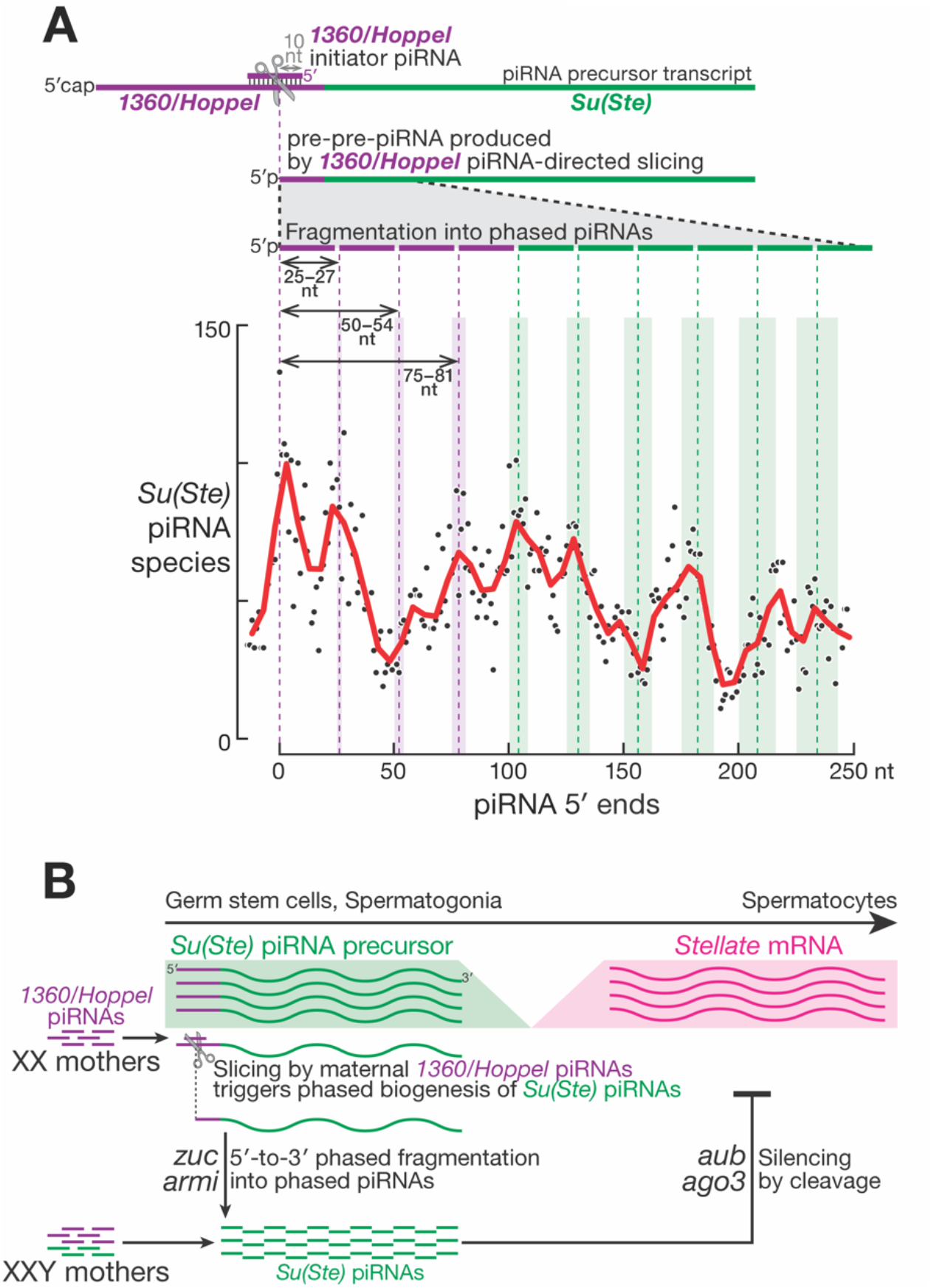
Phased biogenesis of Su(Ste) piRNAs in XXY ovaries and in males. (A) Phased biogenesis of *Su(Ste)* piRNAs in XXY ovaries. (B) Model of developmental regulation of *Su(Ste)* piRNA biogenesis and *Ste* repression in males.

Consistent with Zuc-catalyzed fragmentation of pre-pre-piRNAs into tail-to-head piRNAs, XXY ovaries contained strings of *Su(Ste)* piRNAs in which the 3′ end of one piRNA immediately precedes the 5′ end of another piRNA (Figure S6A; *Zo* = 5.6, *p* = 2 × 10^-8^). Such tail-to-head piRNAs result in the nearly equidistant occurrence of piRNA 5′ ends along a pre-pre-piRNA. Indeed, the 5′ ends of most *Su(Ste)* piRNAs in XXY ovaries concentrated in periodic peaks lying ~26 nt apart starting from *Su(Ste)* pre-pre-piRNA 5′ termini. For example, for *Su(Ste)*-derived long RNAs whose 5′ ends were in the last 100 nt of the *1360/Hoppel* sequence, most *Su(Ste)* piRNA 5′ ends occurred at ~25-27-nt intervals extending as far as ≥ 150 nt into the region of the *Su(Ste)* transcript antisense to *Ste* (Figure 7A, bottom). Thus, the *1360/Hoppel* piRNAs present in XXY ovaries can slice *Su(Ste)* precursors to initiate 5′-to-3′ phased production of *Su(Ste)* piRNAs capable of silencing *Ste*. We note that, because *Ste* loci are co-expressed with *Su(Ste)* in ovaries (Figure S6B), XXY ovaries produce *Ste* piRNAs via ping-pong with abundant *Su(Ste)* piRNAs (Figure S6C, *Z*_10_ = 33.7). Importantly, we did not detect *Ste* piRNAs in XX ovaries, which lack *Su(Ste)* transcripts (Figure 6A).

## Discussion

The piRNA pathway is required for production of functional germ cells in animals. In species like *Drosophila*, whose germline is specified by maternally inherited determinants, the oocyte germ plasm contains piRNA•PIWI complexes that instruct their progeny to silence transposons antisense to the inherited piRNAs. Intergenerational continuity of the piRNA pathway in these species therefore relies on the continued passage of information through the germline. Such maternal inheritance is not possible for Y chromosome-encoded piRNAs, as females lack a Y chromosome. How can mothers instruct their sons to make piRNAs from precursors on the Y chromosome? Our data suggest that the *D. melanogaster* male germline relies on maternally deposited, transposon-derived piRNAs to trigger production of *Su(Ste)* piRNAs antisense to *Ste*. The production of such *Ste*-silencing piRNAs is possible because piRNA-directed cleavage of an RNA triggers the production of tail-to-head strings of piRNA via the phased piRNA biogenesis pathway. This model explains how fly males make piRNAs for which no homologous piRNA guides can be deposited by mothers. Our study also reveals that abundant *Su(Ste)* piRNAs are produced before the onset of transcription of their target, *Ste*. Such spatiotemporal separation may be required for effective repression of *Stellate* mRNAs.

In the fly germline, the proteins Rhino and Kipferl bind heterochromatic piRNA-producing loci and initiate transcription of precursor transcripts from both genomic strands (Klattenhoff et al. 2009; Pane et al. 2011; Mohn et al. 2014; Baumgartner et al. 2022). Promoter-independent, RNA polymerase II transcription of these dual-strand piRNA clusters occurs throughout each locus, ignoring splice sites and polyadenylation sequences (Zhang et al. 2014; Chen et al. 2016; Hur et al. 2016; Andersen et al. 2017). This atypical transcription strategy maximizes production of transposon-targeting piRNAs. *Su(Ste)* piRNA biogenesis in the male germline is unlikely to involve such non-canonical transcription of *Su(Ste).* First, our smFISH experiments detected *Su(Ste)* transcripts from only one genomic strand. Second, loss of *rhi* in fly males has no effect on *Ste* silencing (Chen et al. 2021b).

Taken together, our data suggest that the fly male germline has evolved a strategy that uses maternally supplied, transposon-derived piRNAs to generate Y-chromosome derived, *Su(Ste)* piRNAs that silence the selfish genetic element *Ste*. This strategy allows fly females to instruct their sons to produce piRNAs from sequences absent from the maternal genome. We speculate that this same mechanism may be used by mothers to protect their sons from selfish DNA in other species.

## Acknowledgements

We thank the Bloomington *Drosophila* Stock Center and the Developmental Studies Hybridoma Bank for reagents. We thank Zhao Zhang and Nelson Lau for their helpful discussions, and the Yamashita lab members for comments on the manuscript. The research was supported by the Howard Hughes Medical Institute (YMY, PDZ, SEJ), National Institute of Health (NIH R01 HD109667 to JKK, R35 GM136275 to PDZ, R01GM097363 to AAA and R35 GM130272 to SEJ), and the Whitehead Institute for Biomedical Research (YMY).

## Author contributions

ZGV and YMY conceived the project. ZV, IG, CB and YMY conducted experiments. ZGV, IG, YMY, PDZ designed experiments and interpreted the results. ZGV, IG, MRS, CPC, JKK, TWW, and BWB conducted bioinformatics analysis. PC and AA contributed critical information in the course of the investigation. ZGV, IG, YMY, PDZ wrote and edited the manuscript with the inputs from other authors. YMY and PDZ supervised the research.

## Materials and Methods

### Fly husbandry and strains used

Flies were raised in standard Bloomington medium at 25°C. The following stocks were obtained from the Bloomington Stock Center: *C(1)RM/C(X:Y)y′f′w′, armi^1^, armi^72.1^, aub^HN2^, aub^QC42^, zuc^EY11457^, Df(2L)BSC323, nos-gal4:VP16, bam-gal4:VP16, UAS-gfp-aub, UAS-armi-gfp, UAS-dpp.* RNAi line for *armi:* TRIP.GL00254, *aub:* TRIP.GL00076, *ago3:* TRIP.HMC02938, *vasa:* TRIP.HMS00373, *zuc*: TRIP.GL00111. To generate *UAS-gfp-Ste* (*SteXh:*CG42398), cDNAs was synthetized (Invitrogen, sequence is provided in Supplementary Table S1), and inserted into *UAST-gfp* vector, after the *gfp* cDNA cassette, between BglII and XbaI sites. Transgenic lines carrying these transgenes were generated at BestGene.

To assay male fertility, a single male of indicated genotype (0-1days old) was crossed to three *y^1^w^1118^* virgin females (0-2 days old) at room temperature. Flies were removed after 7 days, and the number of progenies was scored.

### Western blots

Testes (20 pairs/sample) were dissected and rinsed with PBS twice, snap frozen, and kept at −80°C until use. Testes were homogenized in 100μl PBS, supplied with c0mplete protease inhibitor +EDTA (Roche), and mixed with 100μL of 2X Laemmli Sample Buffer (BioRad). Cleared lysates were separated on a 12% Tris-Glycine gel (Thermo Scientific), and transferred onto polyvinylidene fluoride (PVDF) membrane (Immobilon-P, Millipore). The primary antibodies used: mouse anti-α-Tubulin (4.3; 1:3000)(Walsh 1984) obtained from the Developmental Studies Hybridoma Bank), anti-Ste serum (1:10,000). The polyclonal anti-Ste antibody was generated by immunizing guinea pigs with KLH conjugated Ac-KPVIDSSSGLLYGDEKKWC (53-70aa of Ste, Covance, Princeton, NJ). Horseradish peroxidase (HRP)-conjugated goat anti-mouse IgG, and anti-guinea pig IgG (1:10,000; Jackson ImmunoResearch Laboratories) secondary antibodies were used. The signals were detected by Pierce ECL Western Blotting Substrate enhanced chemiluminescence system (Thermo Scientific).

### smRNA-FISH

smRNA-FISH was conducted following the protocol described previously (Fingerhut et al. 2019). DNA oligo probes to detect *Ste* and *Su(Ste)* RNA were conjugated with Quasar 570, Cy3 or Cy5 fluorophores (Biosearch Technologies and IDT, see Supplementary Table S2 for probe information). Testes were mounted using VECTASHIELD media with 4′,6-diamidino-2-phenylindole (DAPI; Vector Labs). Images were captured by a Leica TCS SP8 confocal microscope with a 63×oil-immersion objective (NA = 1.4) and processed by ImageJ software.

### qRT-PCR

Total RNA was isolated by Direct-zol RNA miniprep kit (Zymo Research) from biological triplicates of XY (100 testis/sample), XX or XXY gonads (60 ovary/sample). cDNA was generated by SuperScript III Reverse Transcriptase (Invitrogen) with random hexamer primers. qPCRs of technical triplicates were performed by using Power SYBR Green reagent (Applied Biosystems), and the following primer pairs. *Gapdh:* TAA ATT CGA CTC GAC TCA CGG T and CTC CAC CAC ATA CTC GGC TC, *act5C:* AAG TTG CTG CTC TGG TTG TCG and GCC ACA CGC AGC TCA TTG AG, *Su(Ste):* TTC CGA AGT CAA GCG CTT CAA TG and GGA ATC TGT TTA ATT GCA ACA AC.

Ct values were normalized to *Gapdh* by the ΔΔCt method.

### TaqMan small RNA analysis

The abundance of the following piRNAs were quantified by TaqMan small RNA custom assays (ThermoFisher Scientific): *Su(Ste)-4* piRNA (target sequence: UCU CAU CGU CGU AGA ACA AGC CCG A), the most abundant *Su(Ste)* piRNA (Nagao et al. 2010), *piR-dme-1643 piRNA* (piRBase nomenclature), target sequence: (TAA AGC GTT GTT TTG TGC TAT ACC C), a piRNA we found to be highly abundant in the ovary based on analysis of earlier small RNA sequencing data (Brennecke et al. 2008), and 2S rRNA (target sequence: UGC UUG GAC UAC AUA UGG UUG AGG GUU GUA), which small RNAs we utilized in this study as control. Total RNA was isolated from biological triplicates of XX and XXY ovaries (60/sample) by Direct-zol miniprep kit (Zymo Research). Reverse transcription and qPCR were performed following the manufacturer’s protocol using TaqMan MicroRNA Reverse Transcription Kit, and TaqMan Universal PCR Master Mix II, No UNG (ThermoFisher Scientific). qPCRs were performed in technical triplicates with the appropriate controls. Ct values were normalized to 2S rRNA levels by the ΔΔCt method.

### Small RNA-seq Library Preparation and Analyses

Total RNA from fly ovaries or testis was extracted using the mirVana miRNA isolation kit (Thermo Fisher, AM1560). Small RNA libraries were constructed as described (Gainetdinov et al. 2021) with modifications. Briefly, before library preparation, a spike-in RNA mix, an equimolar mix of six synthetic 5′ phosphorylated RNA oligonucleotides (/phos/UGC UAG UCU UAU CGA CCU CCU CAU AG, /phos/UGC UAG UCU UCG AUA CCU CCU CAU AG, /phos/UGC UAG UCU UGU CAC GAA CCU CAU AG /phos/UGC UAG UUA UCG ACC UUC AUA G, /phos/UGC UAG UUC GAU ACC UUC AUA G, /phos/UGC UAG UUG UCA CGA AUC AUA G), was added to each RNA sample to enable absolute quantification of small RNAs (Supplementary Table S3). To reduce ligation bias and eliminate PCR duplicates, the 3′ and 5′ adaptors both contained nine random nucleotides at their 5′ and 3′ ends, respectively (see below) and 3′ adaptor ligation reactions contained 25% (w/v) PEG-8000 (f.c.). Total RNA was run through a 15% denaturing urea-polyacrylamide gel (National Diagnostics) to isolate 15-29 nt small RNAs and remove the 30-nt 2S rRNA. After overnight elution in 0.4 M NaCl followed by ethanol precipitation, small RNAs were oxidized (to clone only 2′-O-methylated small RNAs) in 40 μl of 200 mM sodium periodate, 30 mM borax, 30 mM boric acid (pH 8.6) at 25°C for 30 min. After ethanol precipitation, small RNAs were ligated to 25 pmol of 3′ DNA adapter with adenylated 5′ and dideoxycytosine-blocked 3′ end (/rApp/NNN GTC NNN TAG NNN TGG AAT TCT CGG GTG CCA AGG/ddC/) in 30 μl of 50 mM Tris-HCl (pH 7.5), 10 mM MgCl_2_, 10 mM DTT, and 25% (w/v) PEG-8000 (NEB) with 600U of T4 Rnl2tr K227Q (homemade) at 16°C overnight. After ethanol precipitation, the 50-90 nt (14-54 nt small RNA+ 36 nt 3′ UMI adapter) 3′ ligated product was purified from a 15% denaturing urea-polyacrylamide gel (National Diagnostics). After overnight elution in 0.4 M NaCl followed by ethanol precipitation, the 3′ ligated product was denatured in 14 μl water at 90°C for 60 sec, 1 μl of 50 μM RT primer (CCT TGG CAC CCG AGA ATT CCA) was added and annealed at 65°C for 5 min to suppress the formation of 5′-adapter:3′-adapter dimers during the next step. The resulting mix was then ligated to a mixed pool of equimolar amount of two 5′ RNA adapters (to increase nucleotide diversity at the 5′ end of the sequencing read: GUU CAG AGU UCU ACA GUC CGA CGA UCN NNC GAN NNU CAN NN and GUU CAG AGU UCU ACA GUC CGA CGA UCN NNA UCN NNA GUN NN) in 20 μl of 50 mM Tris-HCl (pH 7.8), 10 mM MgCl_2_, 10 mM DTT, 1 mM ATP with 20U of T4 RNA ligase (Thermo Fisher, EL0021) at 25°C for 2 h. The ligated product was precipitated with ethanol, and cDNA synthesis was performed in 20 μl at 42°C for 1 hour using AMV reverse transcriptase (NEB, M0277) and 5 μl of the RT reaction was amplified in 25 μl using AccuPrime Pfx DNA polymerase (Thermo Fisher, 12344024; 95°C for 2 min, 15 cycles of: 95°C for 15 sec, 65°C for 30 sec, 68°C for 15 sec; forward primer: AAT GAT ACG GCG ACC ACC GAG ATC TAC ACG TTC AGA GTT CTA CAG TCC GA; reverse primer: CAA GCA GAA GAC GGC ATA CGA GAT XXX XXX GTG ACT GGA GTT CCT TGG CAC CCG AGA ATT CCA, where XXXXXX represents 6-nt sequencing barcode). Finally, the PCR product was purified in a 2% agarose gel. Small RNA-seq libraries samples were sequenced using a NextSeq 550 (Illumina) to obtain 79-nt, single-end reads.

The 3′ adapter (TGG AAT TCT CGG GTG CCA AGG) was removed with fastx toolkit (v0.0.14), PCR duplicates were eliminated as described (Fu et al. 2018), and rRNA matching reads were removed with bowtie (parameter -v 1; v1.0.0) against *D. melanogaster* set in SILVA database (Glockner et al. 2017). Deduplicated and filtered data were analyzed with Tailor (Chou et al. 2015) to account for non-templated tailing of small RNAs. Sequences of synthetic RNA spike-in oligonucleotides were identified allowing no mismatches with using bowtie (parameter -v 0; v1.0.0), and the absolute abundance of small RNAs calculated. The background for *Z*_0_ and *Z*_10_ calculation was all displayed data except positions 0 and 10, respectively.

### Cloning and Sequencing of 5′ Monophosphorylated Long RNAs

Total RNA from fly ovaries or testis was extracted using mirVana miRNA isolation kit (Thermo Fisher, AM1560) and used to prepare a library of 5′ monophosphorylated long RNAs as described (Gainetdinov et al. 2021) with modifications. Briefly, to deplete rRNA, 1 μg total RNA was hybridized in 10 μl to a pool of rRNA antisense oligos (0.05 μM f.c. each) in 10 mM Tris-HCl (pH 7.4), 20 mM NaCl by heating the mixture to 95°C, cooling it at −0.1°C/sec to 22°C, and incubating at 22°C for 5 min. RNase H (10 U; Lucigen, H39500) was added and the mixture incubated at 45°C for 30 min in 20 μl containing 50 mM Tris-HCl (pH 7.4), 100 mM NaCl, and 20 mM MgCl_2_. The reaction volume was adjusted to 50 μl with 1× TURBO DNase buffer (Thermo Fisher, AM2238) and then incubated with 4 U TURBO DNase (Thermo Fisher, AM2238) for 20 min at 37°C. Next, RNA was purified using RNA Clean & Concentrator-5 (Zymo Research, R1016) to retain ≥ 200-nt fragments. RNA was then ligated to a mixed pool of equimolar amount of two 5′ RNA adapters (to increase nucleotide diversity at the 5′ end of the sequencing read: GUU CAG AGU UCU ACA GUC CGA CGA UCN NNC GAN NNU CAN NN and GUU CAG AGU UCU ACA GUC CGA CGA UCN NNA UCN NNA GUN NN) in 20 μl of 50 mM Tris-HCl (pH 7.8), 10 mM MgCl_2_, 10 mM DTT, 1 mM ATP with 60U of High Concentration T4 RNA ligase (NEB, M0437M) at 16°C overnight. The ligated product was isolated using RNA Clean & Concentrator-5 (Zymo Research, R1016) to retain ≥ 200-nt RNAs and reverse transcribed in 25 μl with 50 pmol RT primer (GCA CCC GAG AAT TCC ANN NNN NNN) using SuperScript III (Thermo Fisher, 18080093). After purification with 50 μl Ampure XP beads (Beckman Coulter, A63880), cDNA was PCR amplified using NEBNext High-Fidelity (NEB, M0541; 98°C for 30 sec; 4 cycles of: 98°C for 10 sec, 59°C for 30 sec, 72ºC for 12sec; 6 cycles of: 98°C for 10 sec, 68°C for 10 sec, 72ºC for 12sec; 72°C for 3 min; with the following primers: CTA CAC GTT CAG AGT TCT ACA GTC CGA and GCC TTG GCA CCC GAG AAT TCC A). PCR products between 200-400 bp were isolated with a 1% agarose gel, purified with QIAquick Gel Extraction Kit (Qiagen, 28706), and amplified again with NEBNext High-Fidelity (NEB, M0541; 98°C for 30 sec; 3 cycles of: 98°C for 10 sec, 68°C for 30 sec, 72°C for 14 sec; 6 cycles of: 98°C for 10 sec, 72°C for 14 sec; 72°C for 3 min; forward primer: AAT GAT ACG GCG ACC ACC GAG ATC TAC ACG TTC AGA GTT CTA CAG TCC GA; reverse primer: CAA GCA GAA GAC GGC ATA CGA GAT XXX XXX GTG ACT GGA GTT CCT TGG CAC CCG AGA ATT CCA, where XXXXXX represents 6-nt sequencing barcode). The PCR product was purified in a 1 % agarose gel and sequenced using a NextSeq 550 and NovaSeq (Illumina) to obtain 79+79-nt, paired-end reads.

Sequencing data was aligned to fly genome (dm6) with piPipes (Han et al., 2015b). Briefly, before starting piPipes, sequences were reformatted to remove the degenerate portion of the 5′ adapter (nucleotides 1-15 of read1). The reformatted reads were then aligned to fly rRNA using bowtie2 (v2.2.0). Unaligned reads were mapped to fly genome (dm6) using STAR (v2.3.1), alignments with soft clipping of ends were removed with SAMtools (v1.0.0), and reads with the same 5′ end were merged to represent a single 5′ monophosphorylated RNA species.

## Data Availability

Sequencing data are available from the National Center for Biotechnology Information Small Read Archive using accession number PRJNA879723.

## Notes

### Competing Interest Statement

The authors have declared no competing interest.

