## supplementary information for "*Drosophila* Males Use 5′-to-3′ Phased Biogenesis to Make *Stellate*-silencing piRNAs that Lack Homology to Maternally Deposited piRNA Guides"

#### Supplementary Tables

**Supplementary Table 1: cDNA sequence used for UAS-GFP-Ste construct**

| Gene name | Gene ID | cDNA sequence |
| --- | --- | --- |
| SteXh:CG42398 | 7354447 | ATGTCGAGCTCCCAGAACAACAACAGCAGCTGGATCGATTGGT<br>TCCTCGGGATCAAGGGCAACGAGTTCCTCTGCCGCGTGCCCAC<br>CGACTACGTGCAGGATACGTTCAACCAGATGGGCTTGGAGTAT<br>TTCAGCGAGATACTGGACGTGATCCTGAAGCCGGTGATCGACA<br>GTTCCTCTGGCTTGTTGTACGGCGATGAAAAAAAGTGGTACGG<br>CATGATTCACGCCCGATACATCAAGTCAGAGCGTGGCGTGATT<br>GCTATGCACCGAAAATATATGCGAGGAGATTTTGAATCGTGTC<br>CCAATATCTCCTGTGATAGGCAGAACACCCTGCCAGTGGGCCT<br>CAGCGATGTATGGGGCAAGTCAACCGTCAAGATTTACTGCCCA<br>CGGTGTAAAAAGAACTTTCATCCGAAGTCTGATACACAGCTGG<br>ACGGAGCGATGTTCGGGCCCAGCTTCCCGGACATCTTCTTCTC<br>GCTGCTGCCGAACTTGAGATCGCCCCTGGACGACCCACGTACC<br>TAG |

**Supplementary Table 2: smFISH probes**

|  | DNA oligo sequence | 5' modification | 3' modification |
| --- | --- | --- | --- |
| **ste RNA probe** |  |  |  |
| **ste probe mix** | ccagttcacttgttcacag | - | Quasar 570 |
|  | tctgggagctcgacatgtt | - | Quasar 570 |
|  | ttgatcccgaggaaccaat | - | Quasar 570 |
|  | cggcagaggaactcgttgc | - | Quasar 570 |
|  | tatcctgcacgtagtcggt | - | Quasar 570 |
|  | caagcccatctggttgaac | - | Quasar 570 |
|  | tccagtatctcgctgaaat | - | Quasar 570 |
|  | tcaccggcttcaggatcac | - | Quasar 570 |
|  | caagccagaggaactgtcg | - | Quasar 570 |
|  | tttttttcatcgccgtaca | - | Quasar 570 |
|  | cgtgaatcatgccgtacca | - | Quasar 570 |
|  | ctctgacttgatgtatcgg | - | Quasar 570 |
|  | tgcatagcaatcacgccac | - | Quasar 570 |
|  | ctccttgcatatattttcg | - | Quasar 570 |
|  | ggagatattgggacacgat | - | Quasar 570 |
|  | agggtgttctgcctatcac | - | Quasar 570 |
|  | atcttgacggttgacttgc | - | Quasar 570 |
|  | gctgtgtatcagacttcgg | - | Quasar 570 |
|  | gaagctgggcccgaacatc | - | Quasar 570 |
|  | agcgagaagaagatgtccg | - | Quasar 570 |
|  | caggggcgatctcaagttc | - | Quasar 570 |
|  | cgaaagcctaggtaccgtg | - | Quasar 570 |
|  | tttgggcgatttgagttgc | - | Quasar 570 |
| ***Su(Ste)* RNA probes** |  |  |  |
| *Su(Ste)* probe | attccgaagtcaagcgcttcaat | Cy5 or Cy3 | - |

**Supplementary Table 3: Summary of small RNA-seq library preparation**

| Sample | Replicate | Total RNA, µg | Amount of spike-in, attomol |
| --- | --- | --- | --- |
| XX ovaries (0-5 days old y1w1118 virgin females) | Rep1 | 4.5 | 2000 |
|  | Rep2 | 4.2 | 2000 |
|  | Rep3 | 3.85 | 2000 |
| XXY ovaries (0-5 days old y1w1118/C(X:Y)y'f'w'; nos-gal4/+ virgin females) | Rep1 | 3.65 | 2000 |
|  | Rep2 | 3.4 | 2000 |
|  | Rep3 | 2.8 | 2000 |
| XY testis (0-5 days old y1w1118/Y; nos-gal4:VP16 males) | Rep1 | 6 | 2000 |
|  | Rep2 | 4.9 | 2000 |

#### Supplementary\_Figure\_1

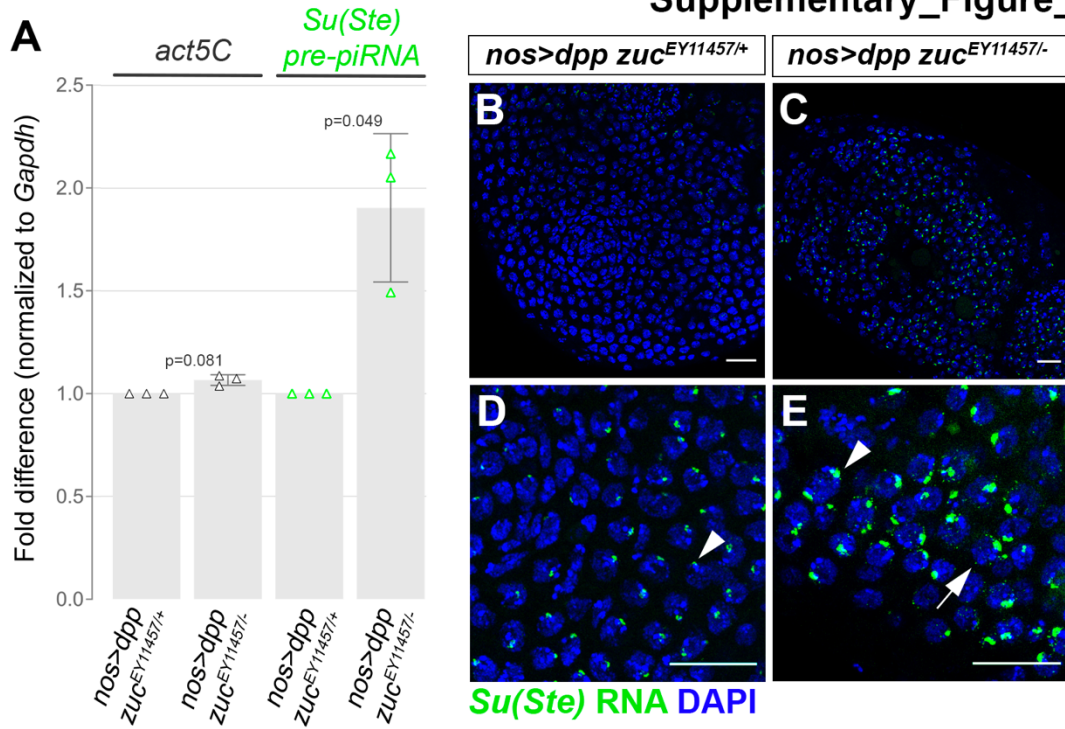

**Supplementary Figure 1. *Su(Ste)* piRNA precursor transcript is not upregulated upon knockdown of *aub*, *vasa*, *ago3*, or *piwi*, Related to Figure 1.** (A) qRT-PCR of *act5C* and *Su(Ste)* precursor RNAs from adult testes of indicated genotypes. Relative abundance normalized to *Gapdh* and heterozygous siblings. SG enriched testes were established by *nos>dpp*). Data are presented as mean±s.d. with p-values of Welch's unequal variances t-tests (unpaired, two-tailed). (B-E) *Su(Ste)* precursor RNAs (green) in adult testes. Arrows point cytoplasmic precursor RNAs, arrowheads point to nuclear transcripts. DAPI (blue), bars 20 μm (B-C) and 5 μm (D-E).

#### Supplementary\_Figure\_2

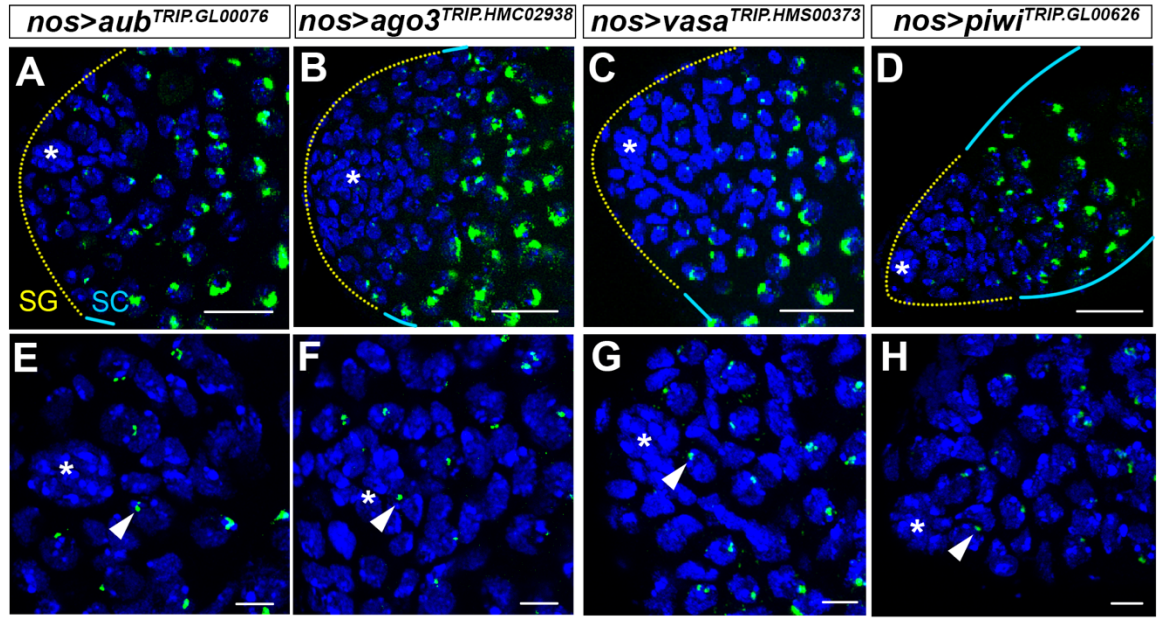

**Supplementary Figure 2. *Su(Ste)* piRNA precursor transcript is not upregulated upon knockdown of *aub*, *vasa*, *ago3*, or *piwi*, Related to Figure 2(A-E)** *Su(Ste)* piRNA precursor transcript (green) in *aub*<sup>RNAi</sup> (A), *ago3*<sup>RNAi</sup> (B), *vasa*<sup>RNAi</sup> (C) and *piwi*<sup>RNAi</sup> (D) testes. (E-H) magnified regions of the niche from A-D. No cytoplasmic foci or enlarged nuclear foci were observed. RNAi constructs were expressed by *nos-gal4*, which drives expression in early germ cells (GSCs and on). The region of GSCs/SGs is indicated by yellow dotted line, SC region by cyan lines. Arrowheads point to nuclear transcripts. Hub (\*), DAPI (blue), bars 20 μm (A-D) and 5 μm (E-H).

##### Supplementary\_Figure\_3

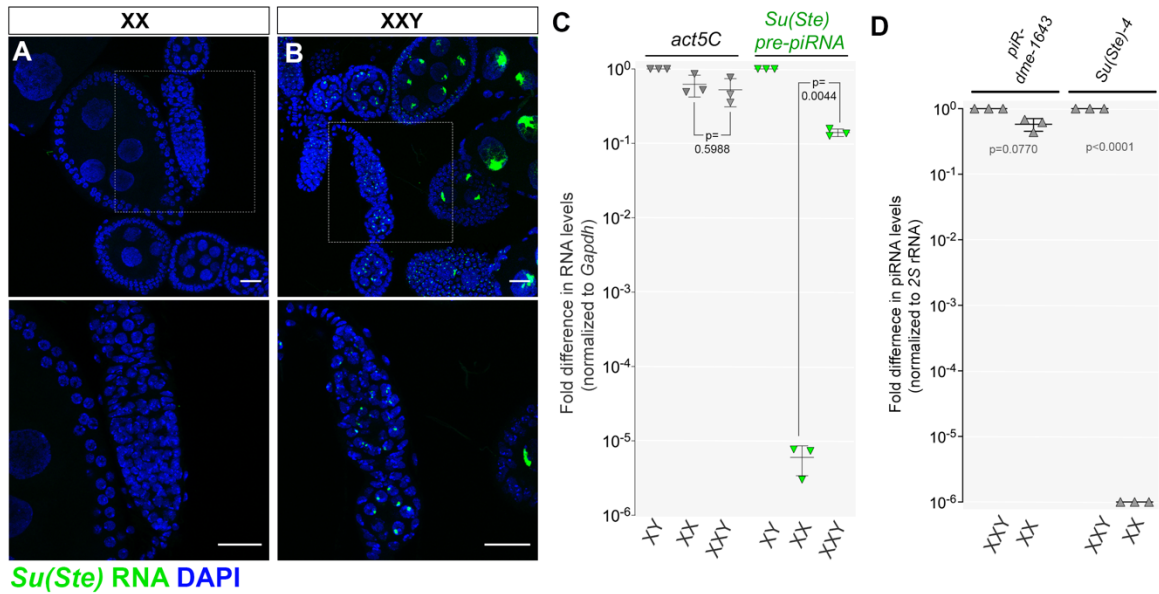

**Supplementary Figure 3. *Su(Ste)* precursor transcripts and piRNAs in XXY ovaries, Related to Figure 6.** (A-B) Germaria and early egg chambers of XX (A) and XXY (B) females with magnified inserts of germaria. *Su(Ste)* piRNA precursor transcript (green), DAPI (blue), bars 20 μm. (C) Relative abundance of *Su(Ste)* piRNA precursor transcript in XY testis, XX ovary vs. XXY ovaries determined by qRT-PCR, normalized to *Gapdh* and XY testes. Data are presented as mean±s.d with p-values of Welch's unequal variances t-tests (unpaired, two-tailed) from biological triplicates. (D) Relative abundance of control (*piR-dme-1643*, *roo* transposon targeting abundant piRNA in ovary) and *Su(Ste)-4* piRNA in XX and XXY ovaries, normalized to 2S rRNA and XXY ovaries. Data are presented as mean±s.d with p-values of Welch's unequal variances t-tests (unpaired, two-tailed) from biological triplicates.

#### Supplementary\_Figure\_4

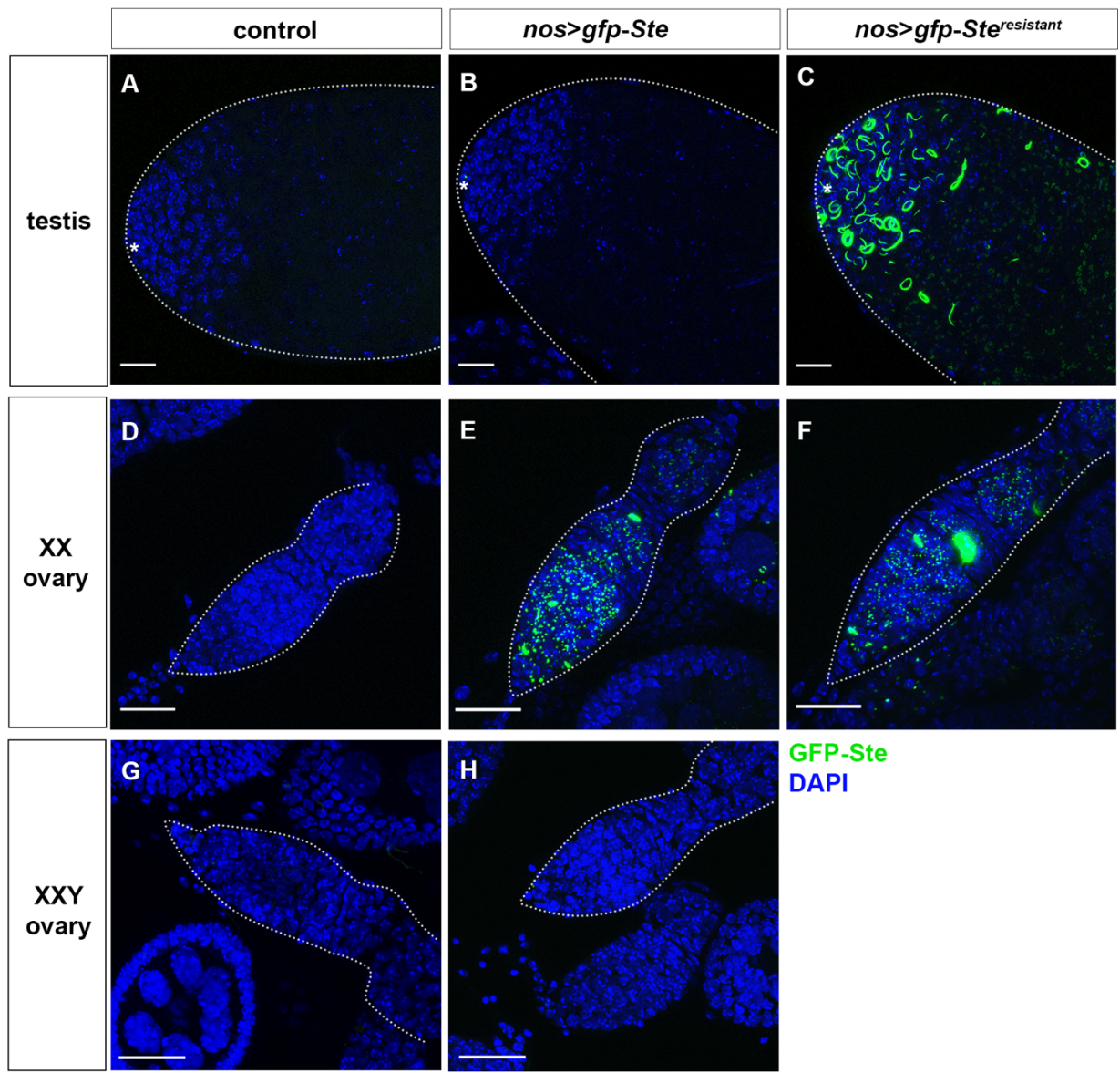

**Supplementary Figure 4. *gfp-Ste* reporter is effectively silenced in the ovary of XXY females, Related to Figure 6.** GFP (green) in testis from XY males (A-C) or germaria from XX (D-F) or XXY (G-H) females. DAPI (blue), bars 20 μm. Asterisks (\*) indicates the hub in A-C.

#### Supplementary\_Figure\_5

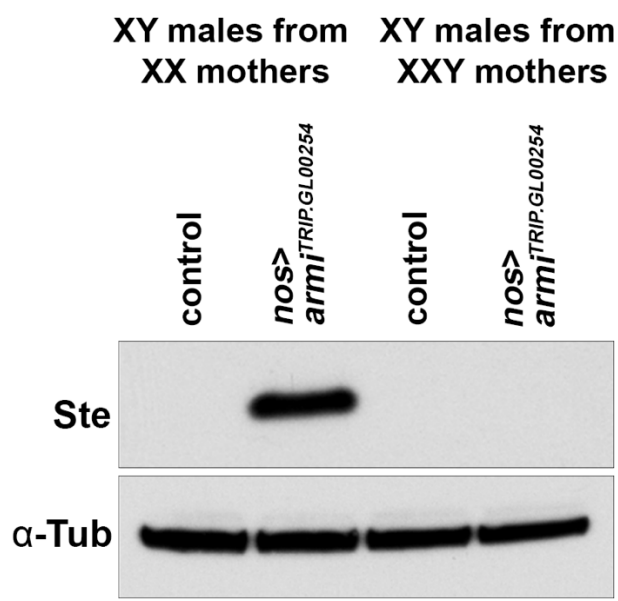

**Supplementary Figure 5. Repression of Stellate protein in *armi<sup>RNAi</sup>* males from XXY mothers, Related to Figure 6.** Anti-Ste and anti-Tubulin Western blotting of testes from indicated genotypes.

### Supplementary\_Figure\_6

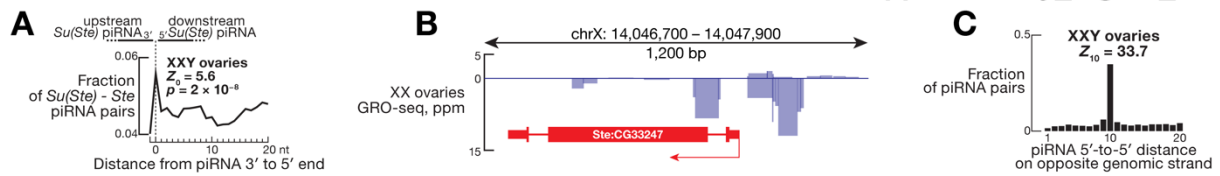

**Supplementary Figure 6. *Su(Ste)* piRNAs make *Ste* piRNAs in XXY ovaries, Related to Figure 7.** (A) Distance between *Su(Ste)* piRNA 5' and 3' ends on the same genomic strand in XXY ovaries. (B) Nascent transcripts (GRO-seq) at a *Ste* locus. Data are from (Wang et al. 2015) for all (uniquely and multiply mapping) reads without apportioning to other *Ste* loci. (C) Ping-pong (10-nt overlap on opposite genomic strands) between *Su(Ste)* and *Ste*-derived piRNAs in XXY ovaries. All data are the mean of three biological samples.

Wang W, Han BW, Tipping C, Ge DT, Zhang Z, Weng Z, Zamore PD. 2015. Slicing and Binding by Ago3 or Aub Trigger Piwi-Bound piRNA Production by Distinct Mechanisms. *Mol Cell* **59**: 819-830.
